# ZDHHC17 Links S-Acylation, Huntington Disease, VCP-associated Multisystem Proteinopathy, and Amyotrophic Lateral Sclerosis

**DOI:** 10.64898/2026.04.05.716148

**Authors:** Firyal Ramzan, Gyana G. Mishra, Yasmeen Alshehabi, Cailyn M. Perry, Lucia M. Q. Jadon, Ashish Kumar, Helen Kim, Ryan Uy, Anthony Dang, Muneera Fayyad, Raoul Ovalekar, Alicia Dubinski, Alyssa E. Johnson, Virginia Kimonis, Christine Vande Velde, Bruce H. Reed, Dale D. O. Martin

## Abstract

Protein mislocalization is an important contributor to neurodegeneration. We have identified disrupted S-acylation as a driver of mislocalization. S11acylation, the addition of long-chain fatty acids to cysteine residues, is mediated by the ZDHHC family of S-acyltransferases, and regulates protein localization by modulating protein hydrophobicity. Among these enzymes, we have identified ZDHHC17 as a central node in neurodegenerative disease. Although decreased ZDHHC17 activity is correlated with Huntington disease (HD) pathology, its broader contributions to neurodegeneration remain poorly defined.

Here, we present new evidence that ZDHHC17 S-acylates or interacts with multiple proteins implicated in amyotrophic lateral sclerosis (ALS), including Valosin-containing protein (VCP) and TAR DNA Binding Protein (TDP1143). We further confirm that VCP, TDP-43, FUS, C9ORF72, and SQSTM1 are S-acylated across rodent models of multiple neurodegenerative diseases, including HD, VCP-associated multisystem proteinopathy, and ALS.

Using biochemical S-acylation assays and confocal microscopy, we show that ZDHHC17 S-acylates VCP, depletes the nuclear localization of both VCP and TDP-43, and modulates VCP-dependent toxicity via ER-stress. We additionally demonstrate that TDP-43 is S-acylated by ZDHHC9.

Motivated by these findings, we investigated the functional significance of motor-neuronal (MN) ZDHHC17 by studying dHip14 (the fly analog of ZDHHC17) in *Drosophila melanogaster*. Here, we found that motor neuron-specific *dHip14* knockdown (KD) in flies resulted in impaired motor function, whereas ubiquitous depletion led to pharate-adult lethality. Together, these results highlight a shared mechanism across neurodegenerative diseases.

Collectively, our findings position ZDHHC17 as a critical enzyme that links fatty acylation, protein homeostasis, and neurodegeneration. Understanding how ZDHHC17 orchestrates S-acylation across different cellular pathways may reveal new therapeutic strategies to restore proteostasis in neurodegenerative disease.

## Introduction

Protein aggregation is a hallmark of nearly all neurodegenerative diseases (Hung & Link, 2011; Suk & Rousseaux, 2020). However, the toxicity of aggregates is a topic of debate within different diseases and is more likely related to the oligomer intermediates. Regardless, aggregation is an indicator of proteostasis deficiency or impaired protein turnover. Importantly, an earlier event that precedes aggregation is protein mislocalization (Ramzan et al., 2023; Suk & Rousseaux, 2020; Wlodarczyk et al., 2024). Mislocalization can simultaneously lead to both a toxic loss of function at a protein’s native site of localization and a toxic gain of function in its new, wrong location. Therefore, understanding the mechanisms of protein mislocalization is critical for identifying early drivers of neurodegeneration.

Protein lipidation, such as fatty acylation, prenylation, and others, are key regulators of protein localization (Aicart-Ramos et al., 2011; Yuan et al., 2024). Of these, S-acylation, commonly known as palmitoylation, is of particular interest as it involves the reversible addition of long-chain fatty acids, typically the 16-carbon palmitate, to specific cysteine residues within proteins via a labile thioester bond (McIlhinney et al., 1985) (for a recent review, see (Chen et al., 2021)). S-acylation is mediated by S-acyltransferases (ZDHHC) enzymes named after their conserved Asp-His-His-Cys (DHHC) active site and zinc (Z) finger domain (Hou et al., 2009; Roth et al., 2006). Typically, S-acylation acts as a molecular “postal code”, directing proteins to different locations within the cell, by manipulating protein hydrophobicity and, thus, affinity for membrane binding (Aicart-Ramos et al., 2011). Therefore, alterations in S-acylation may contribute to protein mislocalization, and ultimately proteostasis deficiencies (Aicart-Ramos et al., 2011; R. Guo et al., 2024; Yanai et al., 2006). Not surprisingly, many ZDHHC enzymes are linked to neurological diseases (Iacoangeli et al., 2020; Jeong et al., 2025; Pinner et al., 2016; Ramzan et al., 2023; Sanders et al., 2016; Singaraja et al., 2011; Wlodarczyk et al., 2024; Yang et al., 2018). Indeed, we previously found that S-acylation is enriched in numerous neurodegenerative diseases, including Huntington Disease (HD), TAR DNA Binding Protein 43 (TDP-43) proteinopathies, and Amyotrophic Lateral Sclerosis (ALS) (Sanders et al., 2015).

Linking many of these diseases, we showed that the autophagy receptor Sequestosome 1 (SQSTM1/p62) is significantly less S-acylated in the brains of HD patients and the YAC128 HD mouse model (Abrar et al., 2025). Because SQSTM1 is critical for autophagy and the removal of misfolded and aggregating proteins, disruptions of SQSTM1 function are linked to many neurodegenerative diseases, including ALS (S. Ma et al., 2019; Martin et al., 2015). Moreover, mutations in *SQSTM1* are associated with ALS and frontotemporal dementia (FTD) (Donáth et al., 2021; Kwok et al., 2014; Layfield & Hocking, 2004; Rea et al., 2014; Yilmaz et al., 2020), suggesting that altered SQSTM1 S-acylation may represent a shared mechanism across diseases.

Protein mislocalization is particularly important in ALS. Aberrant TDP-43 nuclear depletion, and cytoplasmic accumulation, occur in nearly 97% of cases, largely excluding those involving mutations in fused in sarcoma (*FUS)* or superoxide dismutase 1 (*SOD1*) (Hardiman et al., 2017; Kwiatkowski et al., 2009; Mackenzie et al., 2007; Rosen et al., 1993; Vance et al., 2009). Similar to TDP-43, FUS accumulates in the cytoplasm in ALS and FTD patient neurons (Lagier-Tourenne et al., 2010; S.-C. Ling et al., 2013; Portz et al., 2021). A recent study confirmed TDP-43 S-acylation is reduced in SOD1-G93A mice cortices and C9orf72-ALS iPSC-derived motor neurons, regulates its localization in ALS and under proteostasis stress, and consequently impacting the cryptic exon splicing of *STMN2* and *UNC13A* (Xu et al., 2026). Functionally, the loss of nuclear TDP-43 is associated with widespread cryptic exon splicing errors (Bryce-Smith et al., 2025; Fakim et al., 2025; Zeng et al., 2024) and disrupted stress granule dynamics, increasing neuronal susceptibility to stress (Aulas et al., 2012; Aulas & Vande Velde, 2015). Notably, SOD1 is also S-acylated, with disease-associated mutations exhibiting increased S-acylation relative to wild-type protein (Antinone et al., 2013, 2017). Together, these findings prompted us to investigate whether S-acylation serves as a shared mechanism across ALS-associated proteins including SQSTM1, TDP-43, and FUS. As ALS and FTD are highly associated with the Chromosome 9 open reading frame 72 (C9ORF72) hexanucleotide repeat expansions and also exhibit characteristic TDP-43 inclusions and pathology, C9ORF72 was also included (DeJesus-Hernandez et al., 2011; Renton et al., 2011).

Similarly, another rare degenerative disease characterized by impaired proteostasis is valosin-containing protein-associated multisystem proteinopathy (VCP-MSP) (Kimonis et al., 2008; Tresse et al., 2010). VCP-MSP is caused by point mutations in *VCP* (Al-Obeidi et al., 2018; Evangelista et al., 2016; Korb et al., 2021; Meyer & Weihl, 2014; Rea et al., 2014). VCP contributes to many pathways critical for proteostasis, such as endoplasmic reticulum (ER)-associated degradation (ERAD) and autophagy (Chu et al., 2023; Hänzelmann & Schindelin, 2011; Johnson et al., 2015; Meyer & Weihl, 2014; Tresse et al., 2010). Disease-associated VCP variants disrupt these pathways, promoting accumulation of misfolded proteins and cellular stress (Scarian et al., 2022). Indeed, VCP-MSP pathology commonly features SQSTM1 and TDP-43-positive aggregates (Meyer & Weihl, 2014). Given its links to proteostasis, SQSTM1, and TDP-43, altered VCP S-acylation may represent another convergent mechanism across neurodegenerative diseases.

In the current study, we investigate S-acylation as a mechanism of protein mislocalization in neurodegeneration, with a focus on ALS-associated proteins. Herein, we demonstrate that SQSTM1, VCP, TDP-43, FUS, and C9ORF72 are S-acylated in HD, ALS, and VCP-MSP mouse models. We further investigate the S-acyltransferase responsible for modifying these proteins, ZDHHC17. Since S-acylation is a crucial regulator of protein localization and function, our findings demonstrate that disruptions in S-acylation represent a shared mechanism in neurodegeneration.

## Results

## 1. ALS-associated proteins have altered S-acylation in HD

Previously, we found that SQSTM1 S-acylation was significantly decreased in HD patient brains with an approximately 50% reduction and was recapitulated in brains from 8-month-old YAC128 mice, but with a more modest 20% reduction (Abrar et al., 2025). To determine if older mice had a more pronounced reduction in SQSTM1 S-acylation, we measured S-acylation of SQSTM1 in 15-month-old YAC128 mice using the Acyl-Biotin Exchange assay (ABE; Figure 1). Here, we observed a significant decrease of approximately 50% (p<0.05), more in line with previous patient data.

**Figure 1.**
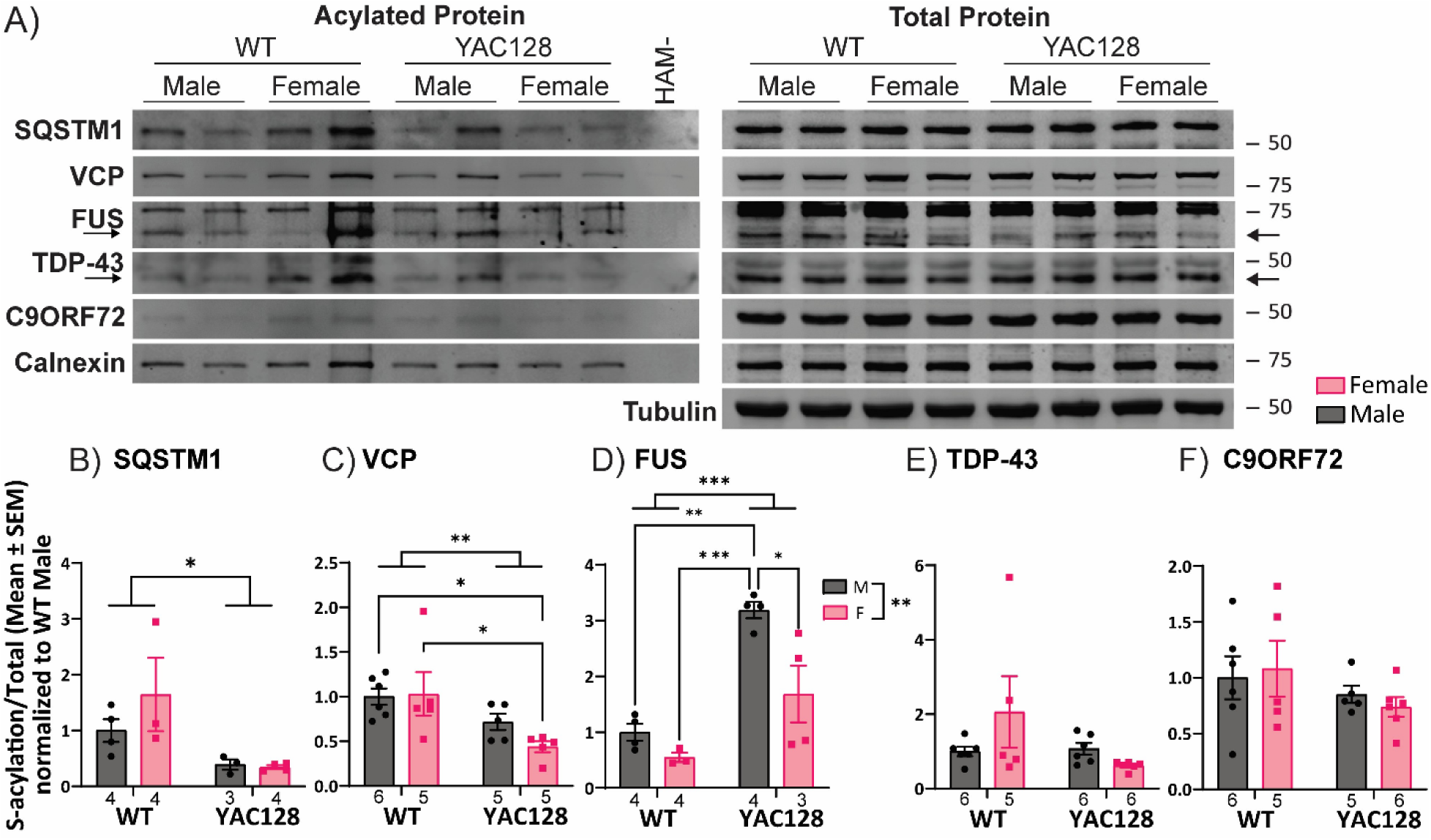
S-Acylation of ALS-associated proteins in YAC128 mouse brains. YAC128 and WT mouse brains demonstrate S-acylation of VCP, TDP-43, SQSTM1, FUS, and C9ORF72. Statistical analysis: A 2-way ANOVA was performed, followed by Tukey’s Multiple Comparisons Test if significant. *p<0.05, **p<0.01, ***p<0.001.

Knowing that SQSTM1 is a direct link between autophagy and proteostasis deficiencies, HD, and ALS, we sought to determine if other ALS-associated proteins were S-acylated. Although more than 40 genes are associated with ALS, the most common mutations occur in a handful of genes, including *TARDBP* (encoding TDP-43), *FUS*, *SOD1*, and *C9ORF72* (Bjelica et al., 2024; Masrori & Van Damme, 2020; Wang et al., 2023). As mentioned, TDP-43 nuclear depletion is a common phenotype in ALS cases. Previous work has demonstrated disease-specific S-acylation of SOD1 (Antinone et al., 2013, 2017). Thus, we sought to investigate whether S-acylation may be a shared feature among ALS proteins and focused on TDP-43, FUS, and C9ORF72. Significantly, VCP’s disease variants are also linked to mislocalization of SQSTM1 (Ju et al., 2009), and several key RNA-binding proteins, including FUS (Harley et al., 2021) and TDP-43 (Hall et al., 2017). Additionally, VCP-associated TDP-43 mislocalization and aggregation occur in multiple tissues, including brain and muscle (Chu et al., 2023), and VCP may have a context-dependent interaction with TDP-43 (G. Liu et al., 2019; Rodriguez-Ortiz et al., 2013), leading us to include VCP in our analysis.

Using ABE, we reconfirmed SQSTM1 S-acylation and demonstrated for the first time TDP-43, VCP, FUS, and C9ORF72 S-acylation in the brains of YAC128 mice, an HD mouse model (Slow et al., 2003). Interestingly, VCP S-acylation was significantly reduced in transgenic mice compared to wild-type littermate controls (p<0.01), and more specifically in YAC128 transgenic females compared to wild-type males (p<0.05) and females (p<0.05) (Figure 1B). Further, FUS S-acylation was substantially increased in the YAC128 mice compared to WT mice (p<0.001), and also increased in males compared to females (p<0.01), and more specifically increased in YAC128 transgenic male mice compared to all other groups (WT-M p<0.01, WT-F p<0.001, YAC128-F p<0.05) (Figure 1C). In contrast, TDP-43 and C9ORF72 were both S-acylated but neither exhibited genotype or sex effects in YAC128 mouse brains. Of note, TDP-43 S-acylation was recently confirmed in ALS models (Xu et al., 2026). Calnexin was included as a stably S-acylated control (Dallavilla et al., 2016).

## 2. ZDHHC17 is a key interactor for proteins associated with disease

As part of an optical screen of TDP-43-GFP and all HA-tagged ZDHHC S-acylating enzymes (data not shown), we noted that TDP-43-GFP mislocalized when co-expressed with HA-ZDHHC9 and HA-ZDHHC17 (Figure 2A). Specifically, TDP-43-GFP is cytoplasmic and depleted from the nucleus with both S-acyltransferases. Interestingly, although both enzymes co-immunoprecipitated with TDP-43-GFP through a GFP-immunoprecipitation (see Figure 2B, HA-ZDHHCs blot) in a click chemistry assay on these samples using alkyne-palmitate (15-HDYA) labelling, only HA-ZDHHC9 appeared to S-acylate TDP-43-GFP (Figure 2B and C, p<0.01). Thus, the two enzymes may lead to TDP-43-GFP mislocalization through distinct mechanisms. Specifically, ZDHHC9 may interact with TDP-43 in an S-acylation-dependent manner, while ZDHHC17 does so independently of TDP-43 S-acylation. This aligns with our previous work, indicating ZDHHC17 interacts with another ALS-linked protein, optineurin (OPTN) in an S-acylation-independent manner (Butland et al., 2014). Furthermore, silencing TDP-43 in SH-SY5Y cells significantly increases *ZDHHC9* and *ZDHHC17* gene expression (Figure 2D, p<0.05; see Supplemental Figure 1 for expression of additional *ZDHHC* genes) suggesting TDP-43 may more broadly regulate total S-acylation. While ZDHHC9 is associated with X-linked disability (Jeong et al., 2025; Masurel Paulet et al., 2014; Shimell et al., 2019), ZDHHC17 has previously known roles in neurodegeneration, particularly HD (Huang et al., 2004, 2011; Martin & Sanders, 2024; Singaraja et al., 2002; Virlogeux et al., 2021; Yanai et al., 2006). Together, these findings suggest ZDHHC17 contributes more broadly to neurodegenerative disease proteins, leading us to investigate ZDHHC17.

**Figure 2.**
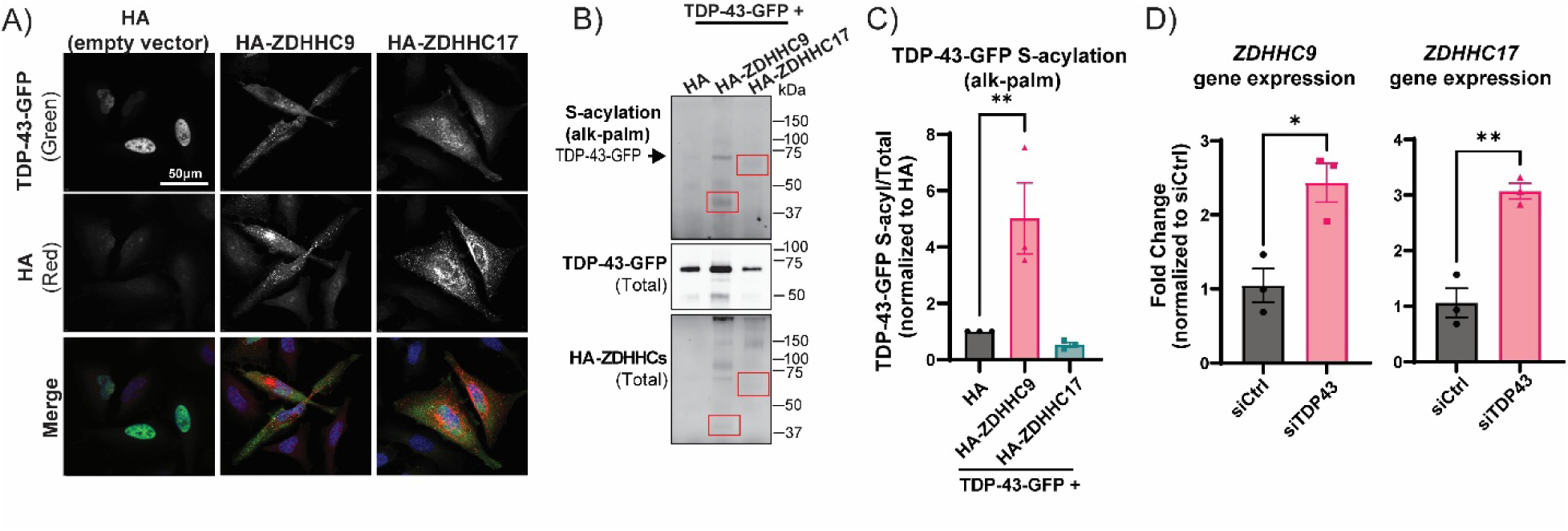
ZDHHC9 and 17 mislocalize TDP-43 while TDP-43 depletion increases ZDHHC expression. **A)** TDP-43-GFP relocalizes from the nucleus to the cytoplasm in the presence of HA-ZDHHC17 and HA-ZDHHC9 in HeLa cells. 3 biological replicates. Western Blot (**B)** and quantifications (**C**) of Click chemistry on GFP immunoprecipitates after incubation with alkyne-palmitate (15-HDYA) demonstrate that HA-ZDHHC9 increases TDP-43-GFP S-acylation. TDP-43-GFP and HA-ZDHHC blots indicate total GFP and HA signal from the TDP-43-GFP immunoprecipitated samples. Red squares indicate locations of HA-ZDHHC9 and HA-ZDHHC17. (**D**) *ZDHHC9* and *ZDHHC17* gene expression upon TDP-43 silencing in SH-SY5Y cells. For additional ZDHHC gene expression data, see Supplemental Figure 1, which also includes *ZDHHC9* and *ZDHHC17* for completeness. Statistical Analysis: (C) One-way ANOVA was performed. Significant results were followed by Fisher’s LSD. (D) Independent t-test were performed. *p<0.05, **p<0.01.

Until recently, there has not been a known S-acylation consensus motif (A. S. Guo et al., 2026) and ZDHHC substrate-binding motifs have been previously identified (Lemonidis et al., 2015; Nadolski & Linder, 2009). We noted a potential ZDHHC17-binding motif within VCP (amino acids ^99^**V**ISI**QP**^104^) (Lemonidis et al., 2015). Conversely, we noted a segment in ZDHHC17’s amino acid sequence a SHP (short-linear binding sequence) box predicted to interact with VCP’s N-domain (^35^YNH**G**Y**G**EP**LG**^49^) (Hänzelmann & Schindelin, 2011, 2016). Based on the observed interaction between TDP-43-GFP and HA-ZDHHC17 (Figure 2B), as well as previous links between VCP and HD (Chu et al., 2023), we sought to confirm if VCP is S-acylated by ZDHHC17. Indeed, VCP-GFP S-acylation is significantly increased in the presence of HA-ZDHHC17 (Figure 3A, p<0.05) as assessed via ABE assay, and the two proteins co-immunoprecipitate (Figure 3B), suggesting that VCP-GFP is S-acylated by HA-ZDHHC17. Additionally, we observed increased cytotoxicity when VCP-GFP and HA-ZDHHC17 were co-expressed. Using a Trypan Blue exclusion assay, we confirmed that VCP-GFP and HA-ZDHHC17 co-expression results in a significant ∼25% reduction in cell viability compared with controls (Figure 3C, p<0.05). Further investigation revealed an increase CHOP, a marker of ER stress, when HA-ZDHHC17 and VCP-GFP are co-expressed, compared with HA-ZDHHC17 alone (Figure 3D, p<0.05), suggesting that this toxicity occurs through the ER stress pathway. Concurrently, when co-expressing VCP-GFP and HA-ZDHHC17, we observed vacuoles reminiscent of paraptosis (Caspase-Independent Cell Death [CICD]; for review see (Hanson et al., 2023)), a non-apoptotic form of cell death mediated by ER stress (Figure 4A, see white arrows).

**Figure 3.**
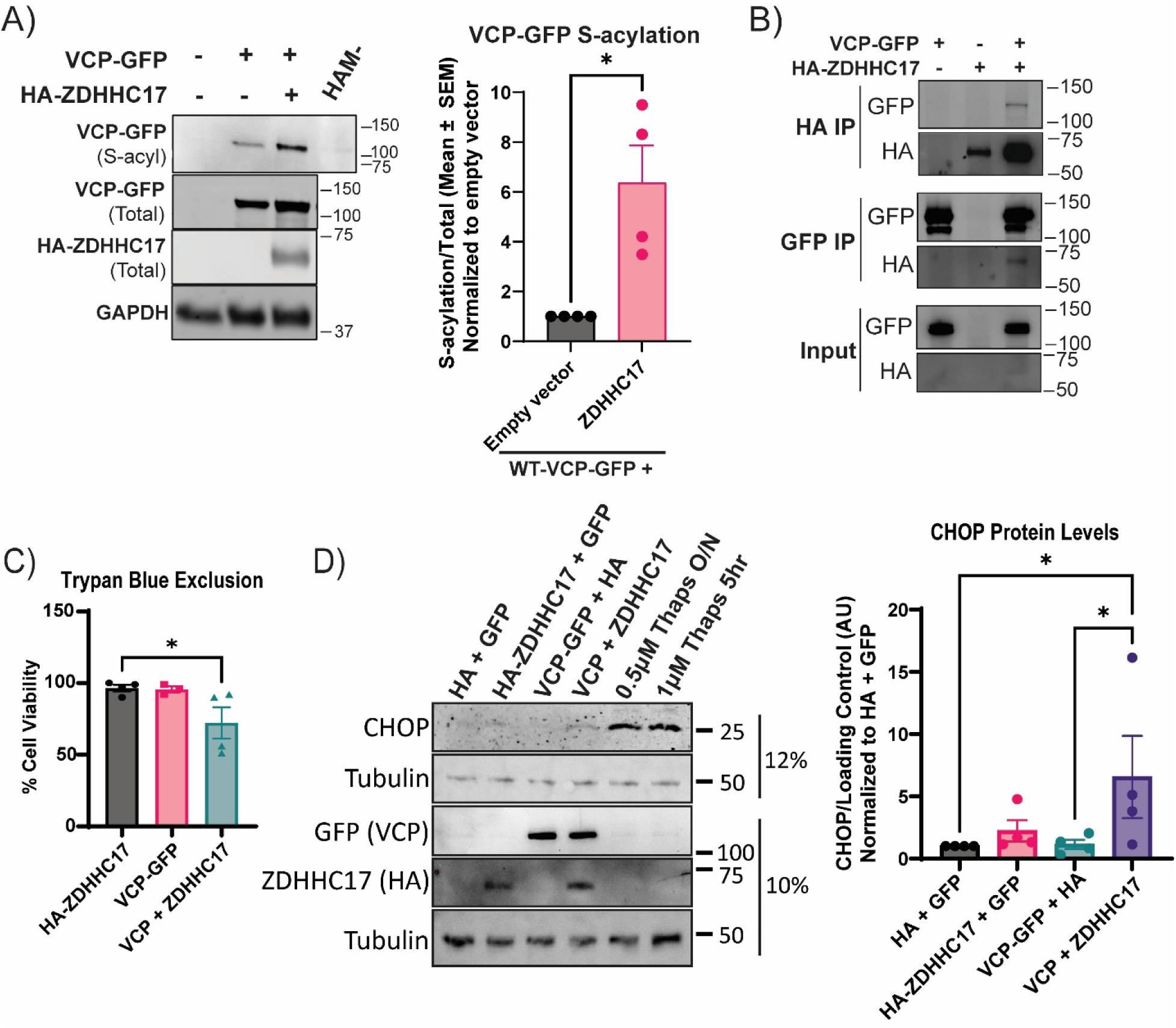
ZDHHC17 interacts with and S-acylates VCP. **A)** ABE results demonstrate that HA-ZDHHC17 increases VCP-GFP S-acylation. Unpaired t-test, *p<0.05. **B)** Co-immunoprecipitation experiments demonstrate that VCP-GFP and HA-ZDHHC17 interact. **C)** Trypan blue exclusion shows reduced % cell viability of HeLa cells co-transfected with HA-ZDHHC17 and WT-VCP-GFP, in comparison to the HA-ZDHHC17 alone control. Significant One-Way ANOVA followed by a t-test, *p<0.05. **D)** Co-expression of WT-VCP-GFP and HA-ZDHHC17 (10% gel) in HeLa cells also results in increased CHOP protein expression (12% gel). Significant One-Way ANOVA followed by Fisher’s LSD, *p<0.05.

**Figure 4.**
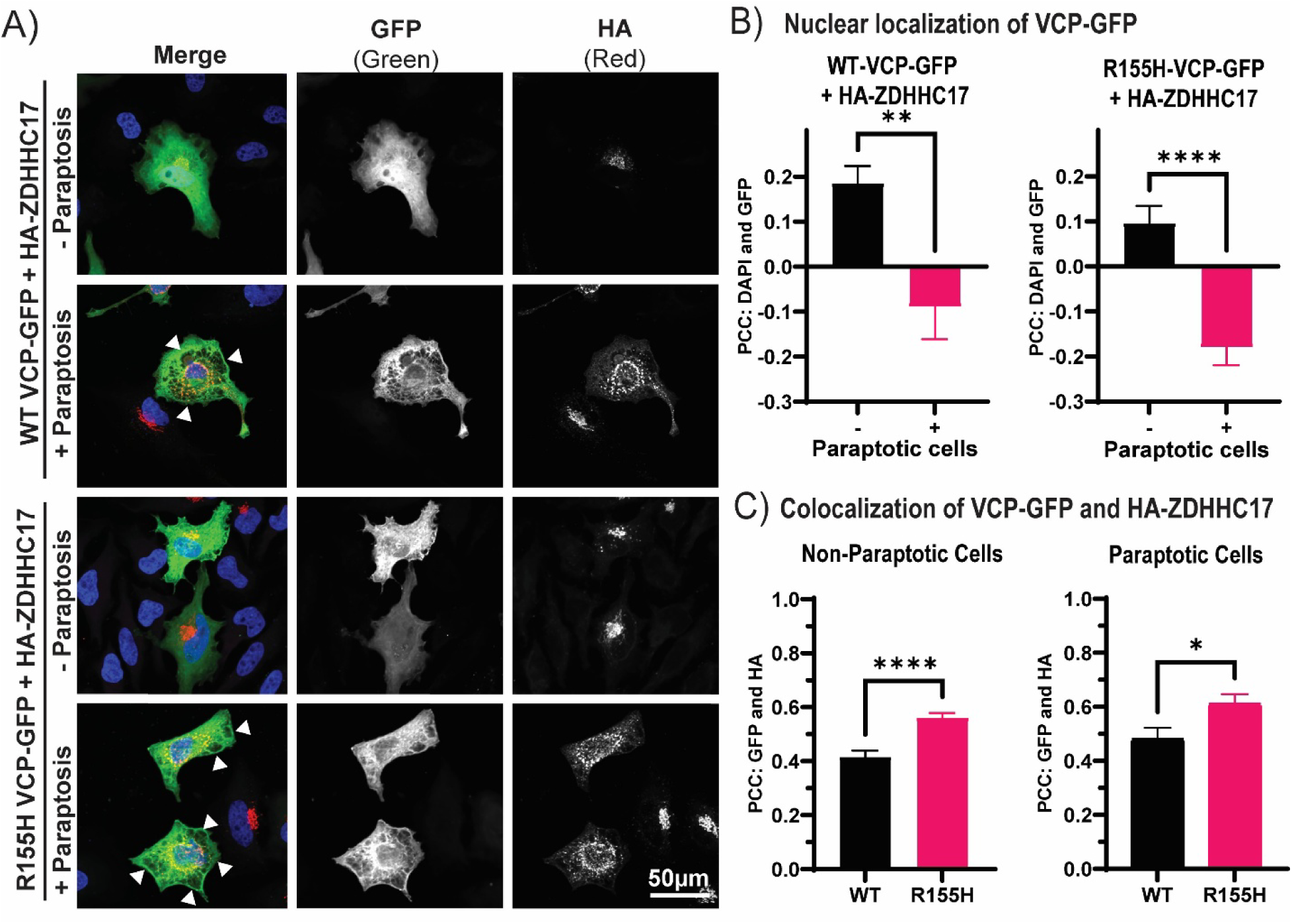
ZDHHC17 colocalization with WT-VCP and R155H-VCP is influenced by paraptosis. **A)** HeLa cells were co-transfected with HA-ZDHHC17 (red, Alexa 594) and VCP-GFP (green). The large cytosolic vacuoles associated with paraptosis are indicated by the white arrows. **B)** Whole cell Pearson’s Correlation Coefficient (PCC) between GFP and DAPI (nuclear indicator) demonstrates that the paraptotic phenotype is associated with increased WT-VCP-GFP and R155H-VCP-GFP nuclear depletion when co-transfected with HA-ZDHHC17. **C)** PCC between GFP and HA in transfected HeLa cells demonstrates that R155H-VCP-GFP has increased co-localization with HA-ZDHHC17 compared to WT-VCP-GFP in both non-paraptotic and paraptotic cells. N = 3, average of 27 cells per condition per replicate. Unpaired T-tests were conducted. *p<0.05, **p<0.01, ****p<0.0001.

Pathogenesis of a disease-associated VCP mutation was recently linked to VCP nuclear depletion (Phan et al., 2024), similar to TDP-43 and FUS. When looking specifically at the paraptotic cells expressing VCP-GFP and HA-ZDHHC17, VCP-GFP nuclear localization is significantly decreased (measured through Pearson Correlation Coefficient (PCC) between GFP and DAPI) (Figure 4B, p<0.01). Among the more than 50 mutations related to VCP-MSP and ALS, R155H-VCP is the most common disease-associated mutation, thus the most studied (Custer et al., 2010; Meyer & Weihl, 2014). Consequently, we looked at nuclear localization of R155H-VCP-GFP. Co-expression of HA-ZDHHC17 with R155H-VCP-GFP reduced nuclear R155H-VCP-GFP, which was also significantly reduced in paraptotic cells (Figure 4B, p<0.0001). Thus, like TDP-43, ZDHHC17 may promote nuclear depletion of WT and R155H-VCP in an S-acylation-dependent manner. We further discovered that R155H-VCP-GFP has increased colocalization with HA-ZDHHC17 compared to WT-VCP-GFP (Figure 4C, p<0.0001), and this is maintained in paraptotic cells (Figure 4C, p<0.05).

Given that VCP S-acylation increased significantly in the presence of HA-ZDHHC17, which was linked to toxicity, we sought to determine whether S-acylation of a VCP disease variant, specifically R155H-VCP, is affected across multiple systems. We found that R155H-VCP-GFP S-acylation significantly increased compared to WT-VCP in transfected HEK293T cells (Figure 5A, p<0.05). Similarly, in a VCP-GFP knock-in model in *Drosophila melanogaster*, S-acylation of the analogous mutation, R152H-VCP-GFP, was significantly higher than that of WT-VCP-GFP (Figure 5B, p<0.05). Importantly, this trend was also observed in patient-derived lymphoblastoid cells, wherein S-acylation of VCP in cells from patients with the R155H-VCP mutation is significantly increased compared to matched unaffected family members (i.e. without a mutation in the *VCP* gene) (Figure 5C, p<0.01). These findings are reinforced by our earlier observation of increased colocalization of HA-ZDHHC17 with R155H-VCP-GFP, which may reflect an increased opportunity for HA-ZDHHC17 to S-acylate R155H-VCP-GFP.

**Figure 5.**
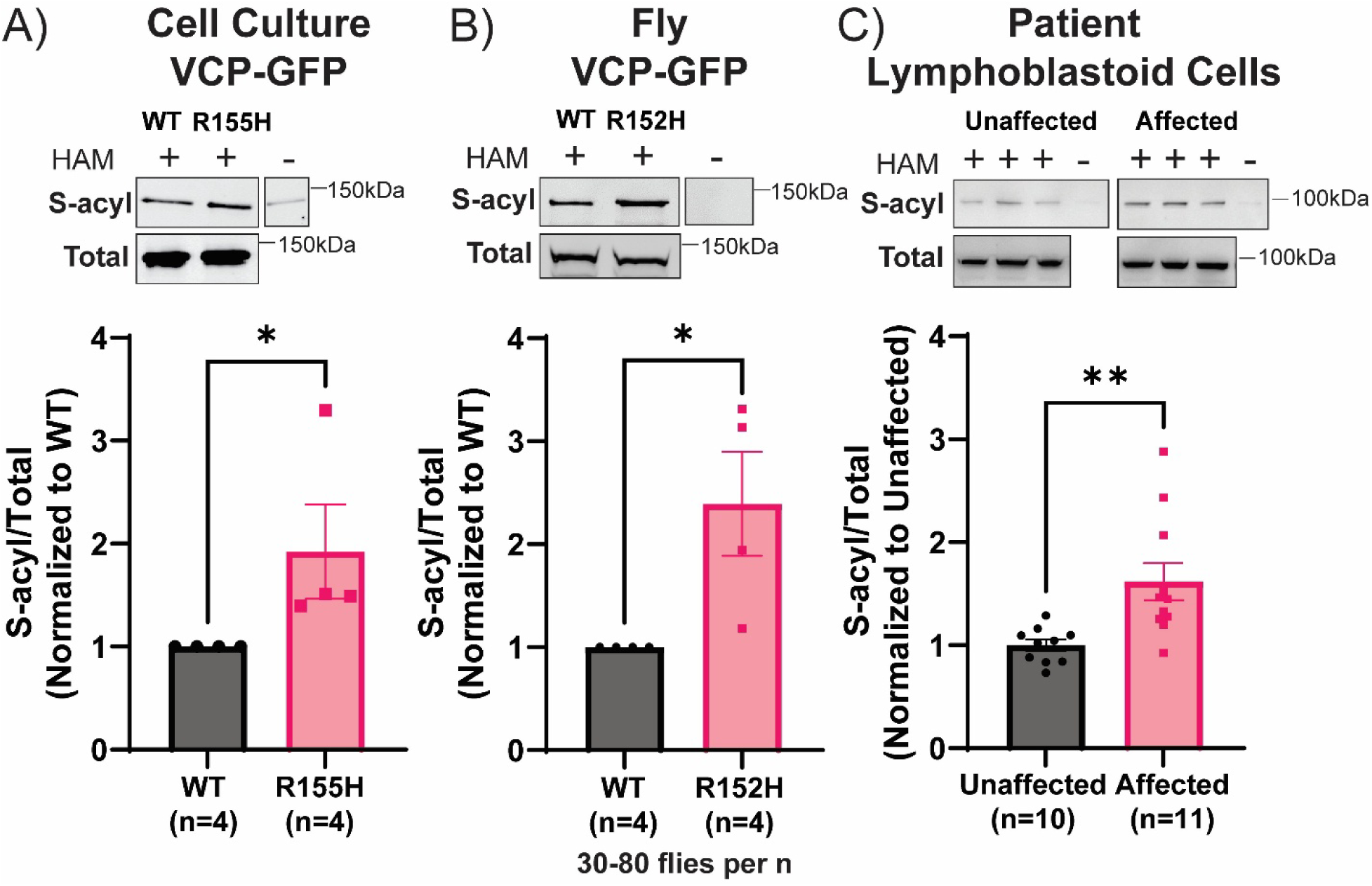
Increased S-Acylation of disease-associated R155H-VCP. S-acylation of disease-associated mutant VCP in **(A)** HEK293T cells (WT-VCP-GFP vs R155H-VCP-GFP), **(B)** *Drosophila melanogaster* flies (WT-VCP-GFP vs R152H-VCP-GFP, R152H corresponds to R155H in human VCP), and **(C)** patient-derived lymphoblastoid cells (Affected = R155H-VCP; Unaffected = no mutation in *VCP* gene). Statistical analysis: Non-parametric Mann-Whitney tests were conducted for transfected cell culture and fly data due to normalization, while an unpaired t-test was run for lymphoblastoid cells. *p<0.05, **p<0.01.

## 3. A fly model of motor neuron-specific *dHip14* (MN-*dHip14*) depletion demonstrates disease-like motor deficits

Given the previously known impact of ZDHHC17 on proteins implicated in neurodegeneration (Huang et al., 2004; Martin & Sanders, 2024; Singaraja et al., 2002; Virlogeux et al., 2021; Yanai et al., 2006) and the alterations in TDP-43 localization, VCP S-acylation, and VCP-associated cytotoxicity, we investigated the functional impact of ZDHHC17 in *Drosophila melanogaster*, which enabled rapid physical and behavioral screening. We crossed flies with a ubiquitous cell promoter (*daGal4*) with *UAS*-*dHip14-RNAi* (*dHip14/*CG6017 is considered the *Drosophila* ortholog of *ZDHHC17*) to produce a fly model of ubiquitous *dHip14 RNAi mediated* depletion (*da*-*dHip14*-KD) (Figure 6A). As validation of the RNAi knockdown, *da*-*dHip14*-KD flies exhibit 40% less *dHip14* expression than controls (Figure 6B, p<0.05). Reduced *dHip14* expression is consistent between early and late pupal stages (Figure 6B). Note, there is no expression data for adult *da*-*dHip14*-KD flies as this global dHip14 knockdown was lethal; development proceeded through the larval-to-pupal transition but resulted in a highly penetrant pharate adult lethal phenotype. (Figures 6D-F). While most *da-dHip14*-KD flies failed to eclose (Figure 6F), some da-*dHip14*-KD flies exhibited lethality during the process of eclosion (Figure 6E). Partially emerged flies were manually removed from their pupal cases prior to lethality and were associated with movement defects (Supplemental Video 1) as well as limb and wing abnormalities (Figure 6E, red and green arrows).

**Figure 6.**
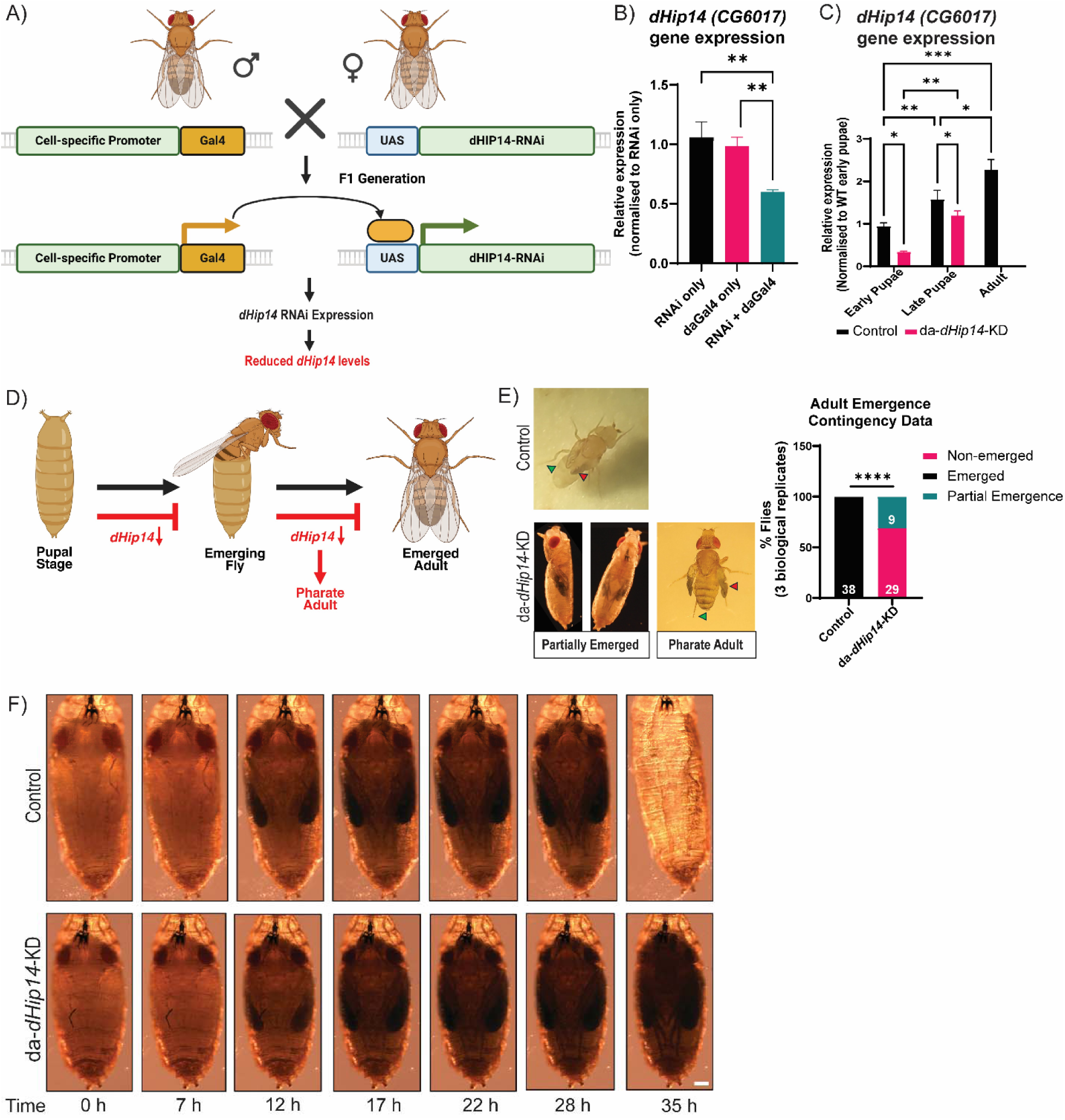
Constitutive ubiquitous *dHip14* (da-*dHip14*) depletion is lethal in flies. **A)** Breeding schematic to produce cell-specific knockdown flies (BioRender). **B)** *dHip14* (*Drosophila* gene CG6017) gene expression in control and *da*-*dHip14*-KD flies. **C)** *dHip14* gene expression across the early pupal, late pupal, and adult stages of development in control and *da*-*dHip14*-KD flies. **D)** Schematic demonstrating fly pupal emergence in black arrows, and pharate adult lethality with ubiquitous *dHip14* depletion in red. Produced in BioRender. **E)** Photographs of a control emerged fly, a partially emerged *da-Gal4*-*dHip14* KD fly, and a *da-Gal4*-*dHip14* KD pharate adult that was dissected from its pupal case. Red and green arrows indicate affected areas of wing and limb development, respectively. Graph indicates the percentage of flies emerging in adulthood and partially (fly numbers in white within the bars). Fisher’s Exact Test for contingencies, ****p<0.0001. **F)** Time-lapse brightfield images of Control and da-Gal4-*dHip14* KD flies over 35 hrs starting at P9-10, scale bar = 200 µm. Control = RNAi only control *p<0.05, **p<0.01, ***p<0.001.

To determine whether the lack of emergence is due to defects in the development of the muscle or the nervous system, we produced flies with pan-neuronal (*Elav-Gal4*), muscle-specific (*mef2-Gal4*), and motor-neuron-specific (*OK6-Gal4*) *dHip14* knockdown. Pan-neuronal RNAi-mediated depletion of dHip14 (*Elav-dHip14*-KD) resulted in complete lethality, while muscle-specific (*mef2*-*dHip14*-KD) and motor neuron-specific (henceforth referred to as MN-*dHip14*-KD) depletion had no impact on viability (Supplemental Figure 2). Climbing assays, which are based on the normal negative geotaxis behavior of wildtype *Drosophila*, were performed on *Mef2*-*dHip14*-KD (not shown) and MN-*dHip14*-KD (Figure 7A). MN-*dHip14*-KD flies exhibit a climbing defect phenotype compared with control flies, with a progressive age-dependent significant decrease (days 7, 14, and 21), indicating that motor neuron-specific *dHip14* depletion potentiates the long-term development of climbing defects in the flies (Figure 7B, sexes combined due to lack of sex differences). Climbing data from *Mef2*-*dHip14*-KD flies suggest a milder impact but is not statistically significant as RNAi-only controls also demonstrate a deficit (Supplemental Figure 3).

**Figure 7.**
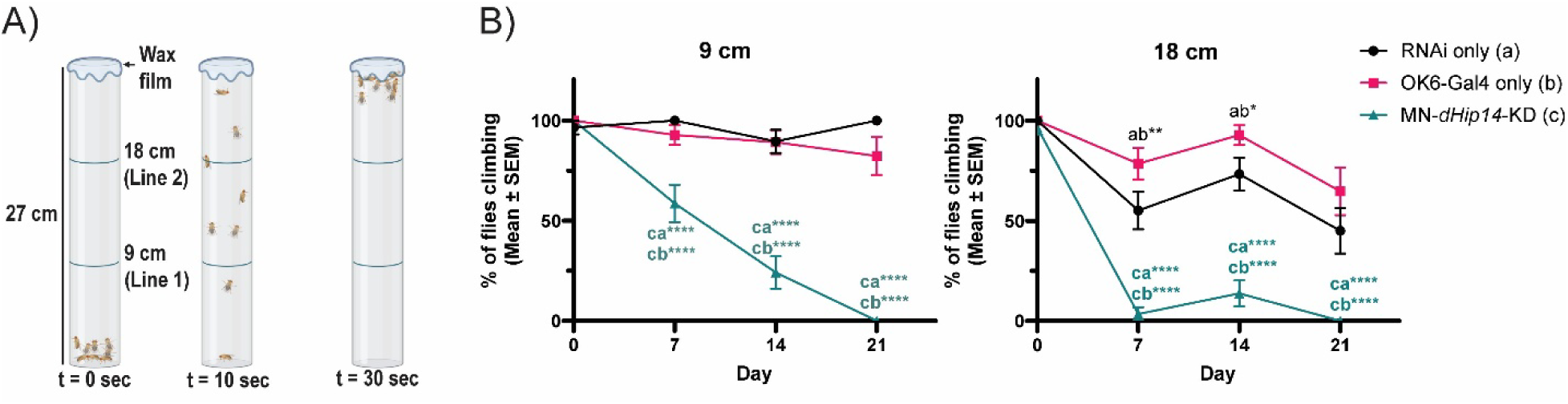
Motor-neuron (MN) specific knockdown of *dHip14*-KD leads to motor deficits in flies. **A)** Diagram demonstrating the negative geotaxis climbing assay (BioRender). Full details in the methods. **B**) Percent of MN-*dHip14*-KD flies climbing to the 9 cm or 18 cm lines over days. 15 Males and 15 females were counted per genotype (3 biological replicates of 5 flies each). No sex effects were found, thus data for both sexes was combined. Statistical analyses: Two-way ANOVAs (Genotype x Day) were run, significant main effects were followed by Fisher’s LSD pairwise analyses. Letters indicate genotype comparisons within day. *p<0.05, **p<0.01, ***p<0.001, ****p<0.0001. See Supplemental Table 1 for a summary of the ANOVA results and pairwise analyses regarding the day factor.

## 4. S-acylation of ALS proteins in VCP-MSP and ALS mouse models

These results prompted us to investigate the S-acylation of VCP and TDP-43 further. Considering the impact of ZDHHC17 on VCP S-acylation and cytotoxicity, as well as the motor degenerative phenotype of MN-*dHip14*-KD flies reminiscent of neurodegenerative disease, we investigated VCP S-acylation in tissues relevant to neurodegeneration in a mouse model of VCP-MSP, specifically in R155H-VCP-KI mice (Badadani et al., 2010), as well as in a mouse model of ALS (M337V-TDP-43 transgenic mice (Gordon et al., 2019)). While the R155H-VCP-KI (R-VCP-KI) mouse model recapitulates several neurodegenerative phenotypes present in patients (i.e. impaired motor behavior, biochemical phenotypes in muscle and bone) (Badadani et al., 2010), it has a mild phenotype, yet is the most suitable mouse model available. This model is complemented by the TDP-43^M337V/M337V^ mouse model, which is reported to recapitulate neurodegenerative phenotypes (Gordon et al., 2019). In the R-VCP-KI mice age-matched to YAC128 mice (Figure 8), we investigated S-acylation of proteins as in YAC128 mice (Figure 1), and observed distinct S-acylation patterns compared with YAC128 mice (Figure 1) and across different tissue types (brain and quadriceps). Specifically, we found that WT and R-VCP-KI mice exhibit genotype-equivalent levels of SQSTM1, VCP, FUS, and TDP-43 S-acylation in the brain (Figure 8A*i-iv*) and quadricep (Figure 8B*i-iv*) tissues, while C9ORF72 did not appear to be S-acylated in either tissue type (Figure 8A and B). In TDP-43^M337V/M337V^ mice, we observed no changes in S-acylation of total TDP-43 (Total, NTg = endogenous TDP-43; TDP-43^M337V/M337V^ = endogenous TDP-43 + hTDP-43-YPet) in the cortex (Figure 9A*i*), but S-acylation was significantly increased in the quadricep of TDP-43^M337V/M337V^ compared to non-transgenic (NTg) controls (Figure 9B*i*), p<0.01). We also observed increased total TDP-43 protein levels in the cortex of TDP-43^M337V/M337V^ mice compared to NTg controls (Figure 9A*iii*, p<0.05). Furthermore, in the quadricep muscle, there was a reduction in endogenous TDP-43 protein in TDP-43^M337V/M337V^ mice compared to hTDP-43^M337V/M337V^-YPet protein levels (Figure 9B*iv*, p<0.01), suggesting that overexpression of mutant TDP-43 results in downregulation of the endogenous protein. In contrast, there is no change in VCP S-acylation across both tissue types.

**Figure 8.**
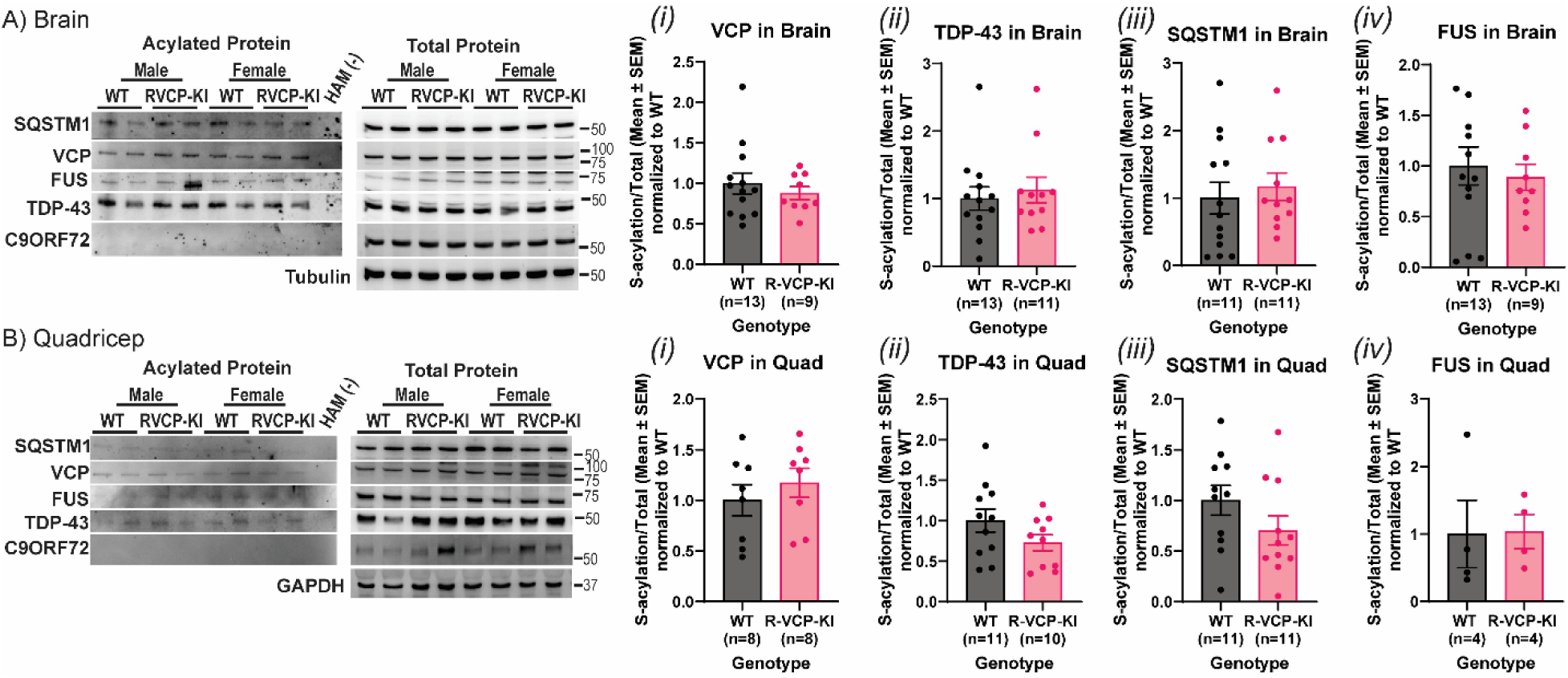
S-acylation of ALS-associated proteins in R155H-VCP Knock-In mouse tissues. R155H-VCP-KI and WT mouse tissues (**A**) brain and (**B**) quadriceps demonstrate S-acylation of SQSTM1, VCP, FUS, and TDP-43. Statistical analysis: Unpaired t-tests were performed.

**Figure 9.**
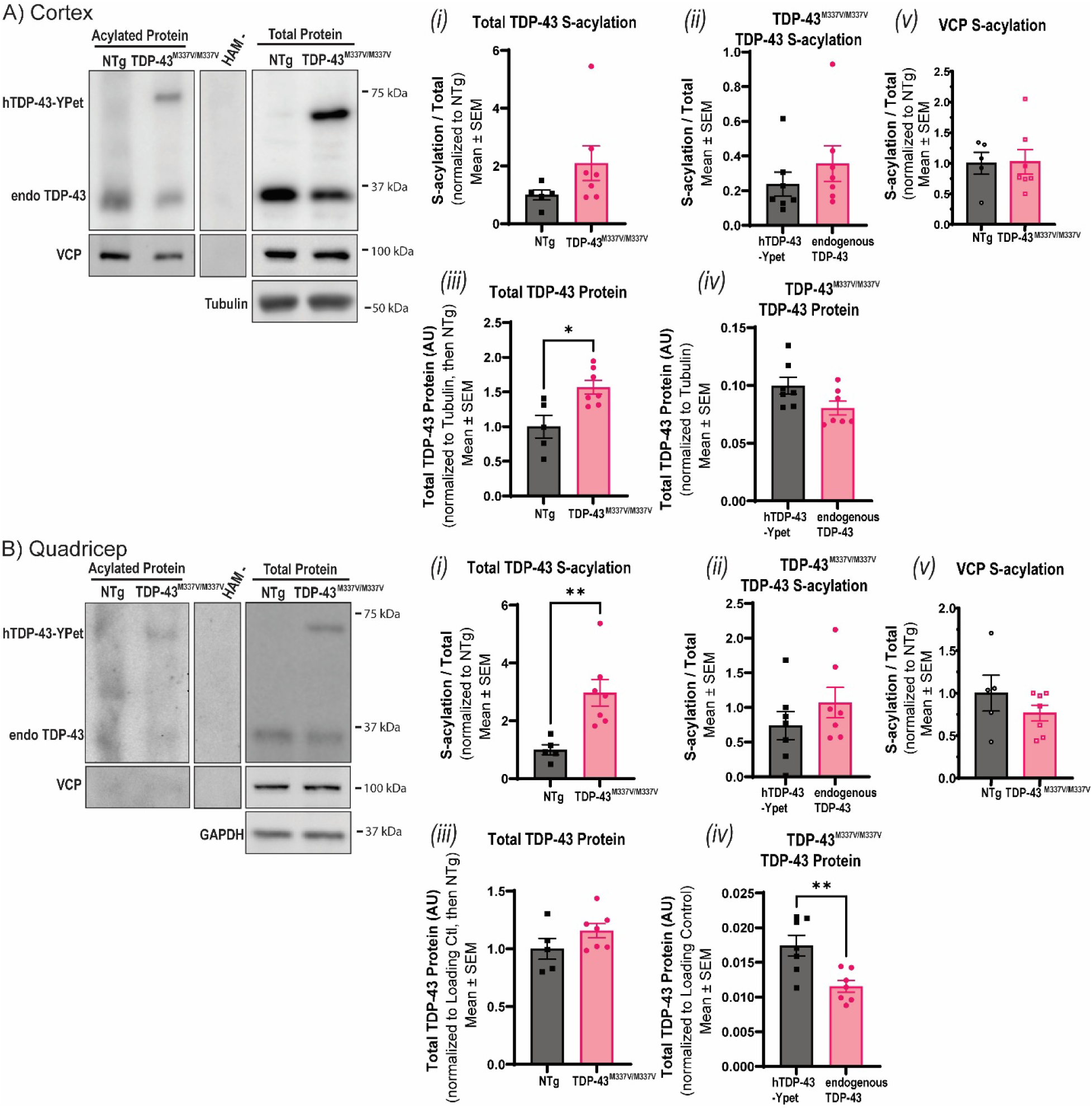
S-acylation of ALS-associated proteins in TDP-43-M337V Transgenic mouse tissues. S-Acylation measured through ABE in TDP-43 M337V transgenic and NTg control mouse tissues **(A)** Cortex and **(B)** quadricep demonstrates S-acylation of TDP-43 and VCP. Total TDP-43 protein levels and TDP-43-M337V S-acylation were also measured. Total TDP-43 in NTg = endo TDP-43, in TDP-43^M337V/M337V^ = endo TDP-43 + hTDP-43-YPet. Unpaired t-tests. *p<0.05, **p<0.01

## Discussion

Herein, we demonstrate that S-acylation of proteins implicated in neurodegeneration is altered in a disease-and sex-specific manner in mouse models of HD and ALS. Of these, we identified, for the first time, that VCP, FUS, and C9ORF72 are S-acylated, and reconfirmed S-acylation of TDP-43 and SQSTM1, revealing S-acylation as a previously unrecognized modification of multiple disease-associated proteins. Notably, TDP-43 is S-acylated and mislocalized by the S-acyltransferase ZDHHC9. In turn, we showed that TDP-43 regulates the expression of multiple ZDHHCs, including ZDHHC9 and ZDHHC17, highlighting a potential negative feedback loop for TDP-43 mislocalization and the potential dysregulation of S-acylation and broader protein mislocalization in ALS.

We further demonstrate that the S-acyltransferase ZDHHC17 plays a role in regulating multiple proteins important in neurodegeneration. Specifically, ZDHHC17 interacts with and causes TDP-43 nuclear depletion independently of TDP-43 S-acylation, whereas it appears to deplete nuclear VCP in an S-acylation-dependent manner. Furthermore, paraptosis, an indicator of cellular stress, is associated with increased nuclear depletion of both WT-VCP and the disease-variant R155H-VCP when co-expressed with ZDHHC17, reminiscent of TDP-43’s nuclear depletion phenotype under cellular stress (Colombrita et al., 2009; Udan-Johns et al., 2014). Consistent with these findings, ZDHHC17 exhibits a toxic interaction with VCP, resulting in reduced cell viability and increased activation of the ER stress pathway, as evidenced by increased CHOP expression. Importantly, ZDHHC17 is critical in normal fly development. Specifically, motor neuron-specific ZDHHC17 plays a critical role in regulating motor ability in flies. Significantly, several ZDHHC enzymes were identified as substrates of TDP-43-regulated cryptic polyadenylation, including ZDHHC17 and ZDHHC24 (Bryce-Smith et al., 2025).

In the HD, VCP-MSP, and ALS disease mouse models, we demonstrate distinct patterns of S-acylation. In the ALS mouse model, we found increased S-acylation of TDP-43 and reduced total TDP-43 protein levels in the quadriceps of TDP-43^M337V/M337V^ transgenic mice compared to non-transgenic controls, suggesting S-acylation in muscles may occur prior to changes in the cortex, though further testing would be needed. TDP-43 has physiological functions in skeletal muscle, including formation of TDP-43-containing myo-granules that regulate local mRNA translation during muscle regeneration (Vogler et al., 2018). Under pathological conditions, these myo-granules can seed amyloid-like TDP-43 aggregates (Vogler et al., 2018), suggesting a potential mechanistic link between typical muscle biology and proteinopathy. Additionally, inclusion body myopathy (IBM), a core disease within VCP-MSP, is characterized by TDP-43 pathology in skeletal muscle ((Kimonis et al., 2008; Meyer & Weihl, 2014)). As such, an increase of TDP-43 S-acylation in quadriceps may reflect an early alteration in TDP-43 regulation within muscle, which may contribute to myo-granule dysfunction and broader proteostasis defects. Future studies examining the relationship between TDP-43 S-acylation, myo-granule formation, and muscle pathology will be important. In contrast, this finding is opposite to recent findings demonstrating reduced TDP-43 S-acylation in brains from an ALS mouse model (SOD1-G93A) and from C9ORF72-iPSC-derived human motor neurons (Xu et al., 2026). Together, these findings may indicate that aberrant S-acylation of TDP-43 may have distinct impacts in distinct tissues, and may ultimately lead to aberrant localization and toxic gain-of-function. Alternatively, mutations in *TDP-43* may lead to increased TDP-43 S-acylation, while mutations in other genes may affect TDP-43 differently, but any changes to TDP-43 S-acylation may lead to mislocalization and proteinopathy. In the HD mouse model, consistent with our previous results in 8-month-old mice (Abrar et al., 2025), SQSTM1 S-acylation was significantly decreased at 15 months. The effect appears greater in older mice (∼20% decrease in 8-month-old mice compared to 50% decrease in 15-month-old mice), which is consistent with mutant HTT S-acylation (Lemarié et al., 2021). Additionally, mutant HTT levels directly correlate with an inverse in ZDHHC17 activity in HD mice and neurons (Huang et al., 2011). As HD mice age, ZDHHC17 activity decreases (Martin & Sanders, 2024; Virlogeux et al., 2021). As another ZDHHC17 substrate, it is not surprising that VCP S-acylation was also significantly decreased in YAC128 mice compared to wild-type controls. Interestingly, FUS S-acylation was significantly increased in YAC128 mice, with female mice showing lower FUS S-acylation compared to male mice. Although findings in the VCP-MSP mouse model did not reach statistical significance, we demonstrated that these proteins are S-acylated in the brain and quadricep tissues. Aging the VCP-MSP mouse model to 15 months may have reduced measurable S-acylation levels, as S-acylation can decrease with age in wild-type and HD mice (R. Guo et al., 2024; Lemarié et al., 2021), likely due to decreased ZDHHC17 activity. Increased S-acylation of VCP’s disease mutant in the cell and fly model systems may indicate higher ZDHHC17 activity compared to the aged VCP-MSP mice. Thus, it will be important to perform age comparisons in the VCP-MSP mice.

Mutations in the *VCP* gene result in VCP-MSP (including FTD, PDB, and IBM) and ALS (Bajc Česnik et al., 2020; Kimonis et al., 2008). We demonstrated that R155H-VCP S-acylation is increased compared to WT-VCP in multiple models, including cells, flies, and patient-derived lymphoblastoid cells. This is supported by increased co-localization of R155H-VCP with ZDHHC17 and increased nuclear depletion of R155H-VCP in paraptosis, indicating a significant role for ZDHHC17 in regulating the function of VCP disease variants.

We have identified ZDHHC17 as a key S-acyltransferase affecting multiple neurodegeneration-relevant proteins. ZDHHC17 was initially identified as a key regulator of huntingtin in HD (Singaraja et al., 2002) and later identified as an S-acylating enzyme (Huang et al., 2004). We have now demonstrated that ZDHHC17 overexpression leads to TDP-43 nuclear depletion without impacting TDP-43 S-acylation, while ZDHHC17 S-acylates and has a toxic interaction with VCP. These findings indicate that ZDHHC17 has a substantial impact on TDP-43 and VCP localization and function. TDP-43’s nuclear depletion in disease is associated with widespread cryptic RNA processing (Brown et al., 2022; Bryce-Smith et al., 2025; Fakim et al., 2025; Klim et al., 2019; J. P. Ling et al., 2015; X. R. Ma et al., 2022; Melamed et al., 2019; Zeng et al., 2024), impaired stress granule dynamics (Aulas et al., 2012; Khalfallah et al., 2018), and impaired DNA damage repair (Mitra et al., 2019). These combined loss-and gain-of-function effects directly impact neuronal susceptibility in neurodegeneration.

VCP’s mislocalization can impact a number of proteostasis pathways. In *Drosophila*, expression of human R155H-VCP disrupts tubular lysosomes and is associated with impaired autophagy dynamics, accumulation of cytoplasmic poly-ubiquitin aggregates, and impaired muscle function (Johnson et al., 2015). Consistent with this, R152H-VCP-GFP in *Drosophila* leads to progressive motor deficits by 4 weeks of age, a shortened lifespan, enlarged SQSTM1 aggregates in 3-week-old adults, and reduced lysosomal network density (Wall et al., 2021). Moreover, the *Drosophila* S-acyltransferase Hip14 is required for lysosome morphology and neuronal integrity in adult brains, functioning as a rate-limiting regulator of lysosomal fusion (Szenci et al., 2025). Herein, we demonstrated that ZDHHC17 has a cytotoxic interaction with WT-VCP, likely driven by ER stress, as indicated by increased CHOP accumulation. These results suggest that ZDHHC17 regulates VCP’s function in ER stress by regulating its S-acylation status. VCP’s functions in ER stress are closely tied to its functions in Golgi assembly and reassembly through the ubiquitin system (Buzuk & Hellerschmied, 2023; Meyer et al., 2012; Shah et al., 2025). Further, under ER stress, VCP is recruited to clear misfolded proteins (Ballar & Fang, 2008); this redirection can delay Golgi reassembly (T. Liao et al., 2024; Shah et al., 2025). This may be linked to Golgi fragmentation observed in ALS and FTD (Haase & Rabouille, 2015; Martínez-Menárguez et al., 2019), both of which are characterized by widespread TDP-43 pathology (reviewed (Scotter et al., 2015)). Given that *VCP* mutations can cause both ALS and FTD phenotypes (Scarian et al., 2022), altered S-acylation-dependent regulation of VCP may represent a common mechanism contributing to degeneration across the ALS-FTD spectrum.

Significantly, we demonstrated that dHip14 depletion in flies results in varying degrees of lethality and motor deficits, depending on the affected tissue. In particular, motor neuron-specific depletion resulted in progressive motor deficits with no impact on lethality, while global and pan-neuronal depletion was lethal. Furthermore, in a mouse model of adult-induced ZDHHC17 knockdown, mutant mice developed progressive, rapid motor deficits, resulting in reduced survival (Sanders et al., 2016). These findings are reminiscent of ALS mouse models, like the C9ORF72 BAC mouse model, which demonstrated progressive and sudden behavioral deficits, particularly in female mutant mice (Y. Liu et al., 2016). Importantly, the C9ORF72 BAC mouse model demonstrated neurodegenerative phenotypes in the motor neurons, which preceded degeneration in the skeletal muscle (Y. Liu et al., 2016). Together, these findings suggest that motor neuron degeneration may have more severe physiological consequences and may precede muscular degeneration. In comparison, overexpression in the TDP-43 and S-acyltransferase screen results in TDP-43 nuclear depletion, a pattern reminiscent of neurodegeneration. As such, alterations in ZDHHC17 levels, whether increased or decreased, may affect the regulation of proteins important in neurodegeneration suggesting ZDHHC17 may play an underappreciated role in regulating ALS proteins.

Our results point to ZDHHC17 as a convergence point for neurodegeneration. ZDHHC17 substrates include proteins involved in trafficking, degradation, and cell stress, as well as many other cellular functions (Butland et al., 2014; Lemonidis et al., 2017; Ohyama et al., 2007; Yanai et al., 2006). Many of these functions, and particularly proteostasis pathways, are commonly disrupted in neurodegeneration, including in VCP-MSP (Badadani et al., 2010; Phan et al., 2024), HD (Lemarié et al., 2021), and ALS (Webster et al., 2017). In particular, synaptic function proteins, including CSPα, SNAP23/25, and MAP6, are interactors of ZDHHC17 (Butland et al., 2014; Lemonidis et al., 2017; Ohyama et al., 2007). An additional factor that may influence disease-specific outcomes is the regional and cellular distribution of S-acyltransferases. For example, the transcriptomic dataset BrainPalmSeq indicates that *ZDHHC17* is broadly expressed across the brain, with higher expression in the frontal and posterior cortices, striatum, and cerebellum, whereas *ZDHHC9* has lower expression (S. Mesquita et al., 2024; Wild et al., 2022). Such variation may influence the relative availability of S-acyltransferases within disease-relevant regions, and future studies investigating ZDHHC gene expression in disease states are needed. A shared reliance on ZDHHC17-regulated S-acylation or interaction may represent a common upstream process that may manifest differently across HD, VCP-MSP, and ALS due to differences in disease-relevant substrates. For example, HTT is the primary driver of pathology in HD, whereas VCP, TDP-43, C9ORF72, and FUS contribute to overlapping pathological mechanisms across ALS, FTD, and VCP-MSP. Notably, TDP-43 mislocalization and aggregation are common pathological hallmarks of numerous diseases, including ALS, FTD, and VCP-MSP (Meyer & Weihl, 2014; Scotter et al., 2015), supporting the hypothesis that dysregulated S-acylation contributes to a broader spectrum of TDP-43 proteinopathies.

## Methods

### Cell Culture and Transfection

HEK293T cells (ATCC; CRL 11268), pre-tested for mycoplasma (https://www.atcc.org/products/crl-11268) or HeLa cells (ECACC; 93021013) were grown at 37°C in 5% CO2 in Dulbecco’s Modified Eagle Medium (DMEM; Wisent # 319-005-CL, supplemented with 10% FBS, 0.1% penicillin–streptomycin, 0.1% L-glutamine, 0.1% sodium pyruvate). HEK293T cells were seeded at 300,000 cells/well (HeLa: 85,000 cells/well) for biochemical experiments or 200,000–225,000 cells/well (HeLa: 75,000 cells/well) in 6-well plates with 0.01 mg/mL poly-D-lysine-coated #1.5 thick coverslips (Fisher Scientific) for microscopy experiments. The next day, cells were transfected with 5 μg of plasmid DNA using calcium phosphate precipitation and incubated for 2 h (or 4 h for HeLa cells), as previously described (Martin et al., 2019). Cells were processed 18 h post-transfection. Plasmids used include N-terminally tagged HA-ZDHHCs (Holland et al., 2016), C-terminally tagged VCP-FEGW and R155H-FEGW plasmids (referred to as WT-VCP-GFP, R155H-VCP-GFP, VCP-GFP, (Ramzan et al., 2024)), and C-terminally tagged pEGFP-TDP-43 plasmid (referred to as TDP-43-GFP, pEGFP-TDP-43 was a gift from Maxime Rousseaux (Addgene plasmid # 235487; http://n2t.net/addgene:235487; RRID:Addgene_235487, (Suk et al., 2025)).

Patient-derived lymphoblastoid cells from affected (R155H-VCP) and unaffected family members (no VCP gene mutation) (Coriell; https://shorturl.at/T9xdm) were grown at 37°C in 5% CO2 in suspension in Roswell Park Memorial Institute Medium (RPMI with L-glutamine and Sodium Bicarbonate; Wisent # 350-000-CL, supplemented with 15% FBS, 0.1% penicillin–streptomycin, 0.1% L-glutamine). At 80% confluency, the cells were collected for biochemical processing.

For S-acyltransferase gene expression studies, SH-SY5Y cells were plated at a density of 5 x 10^5^ cells/well in 6-well plates in Dulbecco’s high glucose modified Eagle medium/Nutrient Mixture F-12 Ham (DMEM-F12, ThermoFisher Scientific) supplemented with 10% FBS (Gibco), 2 mM l-glutamine and 1% MEM non-essential amino acids (ThermoFisher Scientific). 72 h after transfection with 125 pmol Stealth siRNA using Lipofectamine® 2000 (Invitrogen), cells were washed with PBS and processed for RNA extraction. The siRNA sequences used were: Control (siCTL): 5′-AAUUCCAGUUAUUGUAUAUAUUCAG-3′ (Invitrogen), and TDP-43 (siTDP-43): 5′-AAGCAAAGCCAAGAUGAGCCUUUGA-3′ (Invitrogen), as previously described (McDonald et al., 2011).

### Mice

YAC128 mice (FVB/N background, https://www.jax.org/strain/004938# (Slow et al., 2003)) were bred to produce heterozygous or wild-type animals. R155H-VCP knock-in mice (C57B6 background (Badadani et al., 2010)) were bred to produce heterozygous or wild-type animals. Animals were genotyped twice, once using ear notches after weaning and again using tail clips after euthanasia by CO_2_ (as previously described https://www.jax.org/strain/004938<u>#</u>, Badadani et al., 2010). All animals were aged to 15 months prior to euthanasia for tissue collection. All mice were housed in 7.5 × 11.5 × 5 in individually ventilated cages and maintained on a 12 h light cycle starting at 7 am, with ad libitum access to standard mouse chow (8640 Rodent Diet, Envigo) and water. All animal procedures were approved by the University of Waterloo Animal Care Committee and performed in accordance with the Canadian Council on Animal Care (CCAC) guidelines. 10-13-month-old transgenic TDP43^M337V/M337V^ mice express endogenous TDP-43 and two copies of the Ypet-tagged human M337V-TDP43 transgene inserted into the *Rosa26* locus (Gordon et al., 2019). Mice were reared in accordance with protocols approved by the CRCHUM Animal Care Committee and CCAC guidelines.

### Drosophila

All fly cultures were maintained on standard agar-cornmeal-yeast media (11g/L agar, 10g/L cornmeal, 35g/L glucose, 50g/L sucrose, 38g/L Brewer’s yeast, 5g/L yeast extract with trace amount of CaCl_2_ and MgSO_4_ (∼0.1 – 0.5mg/L) and mold inhibitor (Phosphoric acid, propionic acid (∼2mL/L each) and 100mg/L of N-methyl paraben)) at 25°C under a 12 h light/dark cycle. Unless otherwise stated, stocks were obtained from the Bloomington Drosophila Stock Center (BDSC). The Gal4 lines used were pan-neuronal *Elav*-*Gal4* (BDSC 8765, *P{w[+mC]=GAL4-elav.L}2*/*CyO*), motor neuron specific *OK6-Gal4* (BDSC 64199, *P{w[+mW.hs]=GawB}OK6*). The muscle-specific Mef2-Gal4 (BDSC 27390, *P{w[+mC]=GAL4-Mef2.R}3)* and ubiquitous daGal4 (BlBDSC 55850, *If/CyO; P{w[+mW.hs]=GAL4-da.G32}UH1*). Knockdown of *dHip14* was via UAS-RNAi line (BDSC 35012, *P{y[+t7.7] v[+t1.8]=TRiP.HMS01422}attP2*). Heterozygous controls were obtained by crossing to *y w^1118^ P{ry[+t7.2]=neoFRT}19A* (Bloomington Stock 1744; abbreviated as y w^1118^). *R152H-VCP-sfGFP* flies were provided by Dr. Alyssa E. Johnson (Johnson et al., 2015; Wall et al., 2021).

### Negative geotaxis climbing assay

Two lines of heterozygous control flies were obtained by outcrossing respective Gal4 and *UAS-RNAi^dHip14^* strains to y w^1118^. The crosses were carried out on the same day for each biological replicate and grown under a 12 h light:dark cycle at 25°C. F1 progeny that emerged in less than 24 h were collected (to maintain equal age in the cohort) and subjected to the negative geotaxis climbing assay. Briefly, 5 Males and 5 females were collected for each genotype and kept together in a single vial with standard food (10 flies in each vial) to avoid overcrowding. During the climbing assay, flies were anesthetized lightly using CO_2_ and male and female flies were transferred to separate geotaxis tubes (glass tubes measuring ∼27-30 cm). The tubes were kept horizontal for at least 20 mins prior to starting the experiment for acclimatization. Post acclimatization, the geotaxis tubes were tapped gently so that all flies were at the bottom, and the climbing was recorded for 1 min. Number of flies that were able to climb 9 cm and 18 cm mark in 30 secs were recorded. Post assay, the male and female flies were again transferred to a single vial until next timepoint. The assay was conducted on the same day of F1 progeny collection (Day 0) and repeated once every 7 days to assess age-dependent effect on their climbing ability. To avoid any effect due to their circadian cycle, the assay was performed during the same time of the day. The flies were transferred to new vials twice every week depending on the egg laying condition.

### Eclosion Assay

Ubiquitous (*daGal4*) and tissue-specific (pan-neuronal, muscle-specific, and motor neuron-specific) Gal4 driven *dHip14* knockdown crosses, along with their respective y w^1118^ controls, were assessed to evaluate developmental outcomes. Four biological replicates for each cross were used for the assessment. On the first day of pupation, 10 pupae in each vial were circled to ensure accurate tracking. After six days at 23°C under a 12 h light/dark cycle, the vials were inspected, and the total number of flies that successfully eclosed from their pupal cases was recorded. Flies that failed to fully emerge or partially emerged (only the head emerged outside the pupal cases) were identified as pharate adults. Successful eclosion was quantified as the percentage of flies that fully eclosed from their pupae, while pharate counts were recorded in two separate categories (partially emerged and non-emerged).

### Acyl Biotin Exchange (ABE)

The ABE was performed as previously described (Hurst et al., 2017; Martin et al., 2019). Briefly, cells (or 30mg of ground frozen mouse tissue, or 30-60 flies with equal sex distribution) were sonicated in lysis buffer (for cells: 50 mM HEPES, 2% SDS, 1 mM EDTA; for tissues/flies: 10 mM phosphate buffer pH 7.4, 0.32 M sucrose, 1 mM EDTA, 6M Urea with protease inhibitors (PIs)) with 20 mM S-Methyl methanethiosulfonate (MMTS). Lysates were incubated at 50°C for 30 mins to block free thiols with MMTS. After an overnight 80% acetone precipitation at −20°C, protein pellets were washed with 80% acetone and resuspended in 4SB buffer (50mM Tris pH 7.5, 5mM EDTA, 4% SDS with PIs). The resuspension was incubated with 0.7 M hydroxylamine (pH 7.3) and 4 mM HPDP-Biotin (Soltec) for 1 h rotating at room temperature to cleave thioester bonds. After a second overnight 80% acetone precipitation at −20°C, protein pellets were washed with 80% acetone and resuspended in lysis buffer without MMTS. Following protein quantification, 0.5–1 mg of protein was incubated in 150 mM NaCl dilution buffer (50 mM HEPES, 1% TX-100, 1 mM EDTA, 1 mM EGTA) with High-Capacity Neutravidin Agarose beads (Thermo #29202) at 4°C for 4 h. Following two washes with 0.5 M NaCl dilution buffer and one wash with dilution buffer without NaCl, the protein was eluted from beads with elution buffer (0.2% SDS, 250 mM NaCl, 1% β-mercaptoethanol) over a 10 min incubation at 37 °C, then denatured at 95°C for 5 min in 5X sample loading buffer with 5% β-mercaptoethanol, and stored at −20°C.

### Fatty Acid Labelling and Click Chemistry

Cell labelling and click chemistry were performed as described previously (L. M. Q. Liao et al., 2021; Yap et al., 2010). Briefly, cells were deprived of lipids for 45-60 min in DMEM supplemented with delipidated FBS (Life Technologies). Alkyne-tagged fatty acid analog alkyne-palmitate (15-hexadecynoic acid; 15-HDYA; Click Chemistry Tools 1165)) was saponifed in potassium hydroxide. Lipid-deprived cells were then incubated with the saponified alkyne fatty acid analogs for 3 h, after which cells were lysed in modified RIPA buffer (50 mM HEPES pH 7.4, 150 mM NaCl, 0.5% sodium deoxycholate, 2 mM MgCl2, 0.1% SDS, 1% Igepal CA-630). Goat anti-GFP (Eusera) was used to immunoprecipitate (IP) TDP-43-GFP. Immunoprecipitates (IPs) were subjected to click chemistry with tris-carboxyethylphosphine (TCEP), copper sulfate, S-(benzyltriazolylmethyl)amine (TBTA), and azide-647. All reagents for Click Chemistry were purchased from Click Chemistry Tools (Vector Labs).

### Western Blotting

10% or 12% Bis–Tris acrylamide gels were run in XT MOPS buffer (Biorad #1610788) and transferred onto nitrocellulose using the Transblot Turbo (Biorad). Primary antibodies were prepared in 5% skim milk in PBST (0.1% Tween-20 in 1X PBS) and include mouse anti-C9ORF72 (1:1000, Protein Tech (PT) 67824-1-Ig), rabbit anti-Calnexin (1:5000, PT 10427-2-AP), rabbit anti-FUS/TLS (1:2000, PT 68262-1-Ig), rabbit anti-SQSTM1 (1:1000, Invitrogen PA5-20839), mouse anti-VCP (1:1000, PT 60316-1-Ig), rabbit anti-TDP-43 (1:1000, PT 10782-2-AP), rabbit anti-CHOP (1:1000, PT 15204-1-AP), rabbit anti-GFP serum (1:10,000 45min RT, Eusera), and mouse anti-HA (1:1000, 12CA5 clone; Roche 11583816001). Fluorescent secondaries anti-mouse Alexa Fluor 488 (1:2500, Jackson ImmunoResearch 115-545-003) and anti-rabbit Alexa Fluor 790(1:2500, Jackson ImmunoResearch 711-655-152) prepared in 5% bovine serum albumin (BSA) in PBST were used. GAPDH-rhodamine (Biorad), Tubulin-rhodamine (Biorad), and Actin-rhodamine (Biorad) prepared at 1:5000 in 5% BSA in PBST were used as loading controls.

### Fluorescence Microscopy

For fixed cell imaging, HeLa cells were seeded at 75,000 cells/well onto 0.01 mg/mL poly-D-lysine coated coverslips (1.5, VWR) in 6-well plates. After an 18 h transfection (described above), cells were washed once with 1× PBS, then fixed in 4% paraformaldehyde for 20 min at room temperature. After two PBS washes, cells were permeabilized with 0.1% Triton X-100 (with 1 µM MgCl_2_ and CaCl_2_) in PBS for 1 min at room temperature. Following 2 PBS washes, cells were blocked for 30 min with 0.2% gelatin in PBS. Next, coverslips were incubated in primary antibody (rat anti-HA clone 3F10; Roche 12352203 1:500) in blocking buffer in a humidified chamber for 1hr at RT. Following 3 washes with blocking buffer, coverslips were incubated in secondary antibody (rat anti-594 Alex Fluor) in blocking buffer for 1 hr. After 3 washes with blocking buffer, coverslips were incubated in 1 μg/mL DAPI (Sigma D9542) in PBS for 5-20 min at RT, then washed 2 times with blocking buffer. Finally, the coverslips were mounted onto microscope slides (VWR 48311-601) using ProLong Gold antifade mounting media (Invitrogen P36934) and sealed with nail polish. Images were acquired using the Nikon AXR laser scanning confocal microscope at 20× magnification. Using Nikon AXR software, z-stack images at 60x were taken, which were first denoised, then deconvolved, and the resulting maximum intensity projections were used.

### Quantitative PCR (qPCR)

SH-SY5Y cells were washed with PBS and RNA was extracted using the Qiagen RNA Mini Kit. RNA was stored at-80°C until use. cDNA was generated using the High-Capacity cDNA Reverse Transcription Kit (Applied Biosystems, ThermoFisher), and then stored at-20°C until use. qPCR was performed using the PowerTrack SYBR Green Master Mix for qPCR (Applied Biosystems, ThermoFisher) and the QuantStudio 6 Pro thermocycler and analyzed using the Design and Analysis Software 2.0 and Graphpad Prism. Primers used for S-acyltransferases are previously described (McClafferty & Shipston, 2019). The primers for the housekeeping genes are as follows: *ACTB* (fwd: 5’-GCCGCCAGCTCACCA-3’; rev: 5’-CTCGTCGCCCACATAGGAAT-3’), *GAPDH* (fwd: 5’-TCGGAGTCAACGGATTTGGT-3’; rev: 5’-TTCCCGTTCTCAGCCTTGAC-3’). All reactions were run at a melting temperature of 60°C. Target gene expression was normalized to the geometric mean of two housekeeping genes, *ACTB* and *GAPDH* using the delta/delta CT approach.

## Author Contributions

DDOM, FR, GGM, YA, LMQJ contributed to experimental design. YA contributed to data collection for Figures 1 and 2. FR contributed to data collection for Figures 2, 3, 4, 5, and 8. LMQJ contributed to data collection for Figure 2. MF contributed to data collection for Figure 3. AK and RO contributed to data collection for Figure 4. GGM, RU, and HK contributed to data collection for Figures 5, 6, and 7. CMP contributed to data collection for Figures 8 and 9. ADa contributed to microscopy analysis. FR curated and analyzed the data and prepared the figures. GGM curated and analyzed the fly data. FR and DDOM wrote the manuscript. AEJ donated WT-VCP and R152H-VCP *Drosophila* model. CVV and ADu provided RNA for Figure 2D, tissues from M337V-TDP-43 transgenic mice and input on experimental development. VK provided the R155H-VCP-KI mice. BHR provided and maintained the *dHip14-*KD *Drosophila* models and provided input on experimental development. All authors edited the final draft.

## Acknowledgements

We would like to extend our appreciation to HSC, ALS Canada, and Cure VCP Disease, Inc. for supporting this research through funding, discussions, and connections. Thank you to Max Rousseaux for providing TDP-43-GFP plasmids and input.

We would like to acknowledge that the University of Waterloo resides on the traditional territory of five Indigenous communities, including the Ho-de-no-sau-nee-ga (Haudenosaunee), Mississaugas of the Credit First Nation, Anishinabewaki, Attiwonderonk (Neutral), and Mississauga peoples, which is situated on the Haldimand Tract, a land granted to the Six Nations, encompassing six miles on either side of the Grand River. This land is home to many past, present, and future Indigenous peoples.

