## Supplemental Table 1 for "ZDHHC17 Links S-Acylation, Huntington Disease, VCP-associated Multisystem Proteinopathy, and Amyotrophic Lateral Sclerosis"

**Supplemental Table 1. MN-*dHip14*-KD 2-way ANOVA results and pairwise analyses of the Day factor.**

| Comparison | | | | |
|---|---|---|---|---|
| **9cm (Combined Sexes) 2-Way ANOVA** | | | | |
| | **F** | **df** | **p-value** | |
| *Day* | 28.61 | 3, 302 | <0.0001 | |
| *Genotype* | 104.3 | 2, 302 | <0.0001 | |
| *Interaction* | 17.67 | 6, 302 | <0.0001 | |
| **9cm (Combined Sexes) Follow-up analyses using Fisher's LSD - Days only** | | | | |
| RNAi only (a) | **t** | **df** | **p-value** | **Effect** |
| *0 vs. 7* | 0.4757 | 302 | 0.6346 | |
| *0 vs. 14* | 0.9515 | 302 | 0.3421 | |
| *0 vs. 21* | 0.4233 | 302 | 0.6724 | |
| *7 vs. 14* | 1.427 | 302 | 0.1546 | |
| *7 vs. 21* | 0 | 302 | >0.9999 | |
| *14 vs. 21* | 1.27 | 302 | 0.2051 | |
| OK6-Gal4 only (b) | | | | |
| *0 vs. 7* | 0.9768 | 302 | 0.3295 | |
| *0 vs. 14* | 1.465 | 302 | 0.1439 | |
| *0 vs. 21* | 2.093 | 302 | 0.0372 | 0 > 21 |
| *7 vs. 14* | 0.4842 | 302 | 0.6286 | |
| *7 vs. 21* | 1.238 | 302 | 0.2168 | |
| *14 vs. 21* | 0.8169 | 302 | 0.4146 | |
| RNAi + OK6 Gal4 (c) | | | | |
| *0 vs. 7* | 5.709 | 302 | <0.0001 | 0 > 7 |
| *0 vs. 14* | 10.47 | 302 | <0.0001 | 0 > 14 |
| *0 vs. 21* | 12.28 | 302 | <0.0001 | 0 > 21 |
| *7 vs. 14* | 4.757 | 302 | <0.0001 | 7 > 14 |
| *7 vs. 21* | 7.196 | 302 | <0.0001 | 7 > 21 |
| *14 vs. 21* | 2.963 | 302 | 0.0033 | 14 > 21 |
| **18cm (Combined Sexes) 2-Way ANOVA** | | | | |
| | **F** | **df** | **p-value** | |
| *Day* | 53.08 | 3, 304 | <0.0001 | |
| *Genotype* | 74.39 | 2, 304 | <0.0001 | |
| *Interaction* | 8.518 | 6, 304 | <0.0001 | |
| **18cm (Combined Sexes) Follow-up analyses using Fisher's LSD - Days only** | | | | |
| RNAi only (a) | **t** | **df** | **p-value** | **Effect** |
| *0 vs. 7* | 5.118 | 304 | <0.0001 | 0 > 7 |
| *0 vs. 14* | 3.07 | 304 | 0.0023 | 0> 14 |
| *0 vs. 21* | 5.673 | 304 | <0.0001 | 0 > 21 |
| *7 vs. 14* | 2.091 | 304 | 0.0374 | 7 < 14 |
| *7 vs. 21* | 1.049 | 304 | 0.2949 | |

| | | | | |
|---|---|---|---|---|
| *14 vs. 21* | 2.943 | 304 | 0.0035 | 14 > 21 |
| OK6-Gal4 only (b) | | | | |
| *0 vs. 7* | 2.425 | 304 | 0.0159 | 0 > 7 |
| *0 vs. 14* | 0.8083 | 304 | 0.4196 | |
| *0 vs. 21* | 3.464 | 304 | 0.0006 | 0 > 21 |
| *7 vs. 14* | 1.603 | 304 | 0.1101 | |
| *7 vs. 21* | 1.352 | 304 | 0.1774 | |
| *14 vs. 21* | 2.745 | 304 | 0.0064 | 14 > 21 |
| RNAi + OK6 Gal4 (c) | | | | |
| *0 vs. 7* | 10.63 | 304 | <0.0001 | 0 > 7 |
| *0 vs. 14* | 9.448 | 304 | <0.0001 | 0 > 14 |
| *0 vs. 21* | 9.807 | 304 | <0.0001 | 0 > 21 |
| *7 vs. 14* | 1.181 | 304 | 0.2385 | |
| *7 vs. 21* | 0.3503 | 304 | 0.7264 | |
| *14 vs. 21* | 1.401 | 304 | 0.1622 | |
